# A cross-week analysis of urinary extracellular vesicles after respiratory tract exposure intervention identifies systemic signaling changes – a pilot study of isocyanate-exposed workers

**DOI:** 10.1101/2025.02.21.639517

**Authors:** Ikjot S. Sohal, Anoop Pal, Jack Lepine, Pengyuan Liu, Adam Wisneski, Carrie Redlich, Dhimiter Bello

## Abstract

Extracellular vesicles (EVs) from various biofluids have demonstrated potential as prognostic and diagnostic biomarkers in several diseases. However, their utility in early stages of disease development has not been extensively investigated. Additionally, given their role in intercellular communication, decoding the landscape of underlying cell-to-cell signaling in a clinical or subclinical condition based on EV-specific analyses has so far remained challenging. Here, we demonstrate the utility of urinary EVs for cross-week biomonitoring and for capturing the perturbation of underlying organismal signaling in a small well-characterized group of workers with inhalational exposure to methylene diphenyl diisocyanate (MDI) in fabric coating factory. Workers wore organic vapor cartridge respirators during the week to eliminate respiratory exposures. Spot urine was collected before and after the respirator intervention – pre-RESP and post-RESP, respectively. Following extensive characterization of EVs isolated from the urine samples, the relative enrichment of EV-specific RNAs and proteins and their associated biological processes were determined between pre-RESP and post-RESP samples. Distinct EV-specific RNA and protein signatures between the two groups strongly correlated with established biological and immune-related processes involved in asthma and lung inflammation. Using single cell transcriptomics data from the human cell atlas, EV-mRNAs highly enriched in pre-RESP, but not post-RESP, samples revealed global immune dysregulation and enrichment of associated cell types, such as monocytes, macrophages, and neutrophils. The findings indicate that urinary EVs and their contents can be utilized for biomonitoring of lung health and can reveal the effects of exposure on the underlying organismal signaling.

**One Sentence Summary:** Urinary extracellular vesicles and their cargo hold great potential to monitor lung health and changes in systemic signaling in inhalational exposures.

## INTRODUCTION

The human body relies on coordinated cell-to-cell signaling to regulate homeostasis and disease progression. One of the emerging mediators of intercellular signaling are extracellular vesicles (EVs) (*1*). EVs are a group of heterogenous membrane bound vesicles that contain biologically active molecules such as lipids, DNAs, RNAs, and proteins (*2*, *3*). As intercellular communication mediators, the secretion and content of EVs dynamically changes during homeostasis and disease progression, providing crucial molecular information regarding the health status of a certain tissue, organ, or individual. Indeed, EVs isolated from the blood and urine, and their cargo, have been proposed as potential biomarkers for the diagnosis and prognosis of various types of cancers, neurodegenerative, renal, and diabetic diseases (*4–19*).

A prevailing approach for biomarker discovery involves comparing two distinct patient groups, such as healthy vs. diseased individuals or placebo vs. therapy group, which only provides a static view of the disease progression or resolution. However, progression or resolution of a disease is more dynamic. In addition, recent studies have shown high biological inter-individual variation in circulating EVs across healthy humans (*20*). Therefore, developing robust approaches to use EV-based liquid biopsies to capture dynamic changes in disease progression or resolution and changes in the underlying systemic signaling represents a huge opportunity to personalize interventions and treatments, as well as to understand temporal toxicant exposure-disease relationships. As an example, longitudinal analysis of plasma EVs from same individuals has been shown to allow monitoring of disease progression and disease resolution upon therapeutic intervention (*21*). However, frequent blood draws are invasive and can be difficult to obtain under certain scenarios. Urine, on the other hand, is a non-invasive biological medium that is particularly easy to collect in a variety of settings and human subjects. Urinary EVs, however, have been largely studied in the context of urogenital tract-related health and disease. Here, we demonstrate the utility of urinary EVs for probing lung health and for capturing the perturbation of underlying systemic cellular signaling across one week of respiratory exposure intervention in a group of workers with inhalational exposure to methylene diphenyl diisocyanate (MDI) in a fabric coating factory.

A group of 5 healthy workers were selected with well-characterized chronic occupational exposure to methylene diphenyl diisocyanate (MDI) – a potent sensitizer and asthmagen – during manufacturing of polyurethane fabrics. Urine samples were collected as part of a one-week intervention study, which required the workers to wear half-facepiece organic vapor cartridge (OVC) respirators to eliminate inhalation exposure. The urinary EVs were isolated from urine samples collected before the respirators were worn (i.e. at the beginning of work week) and at the end of the work week, referred to here as pre-respirator intervention (pre-RESP) and post-RESP, correspondingly. The isolated EVs were extensively characterized for their morphology, purity, size distribution, and concentration. Upon profiling of the EV cargo for global changes, we observed marked changes in urinary EV-mRNA, -ncRNA (non-coding RNA), and -proteome of these workers after one week of respirator intervention. The different EV cargo types consistently correlated with biological processes associated with asthma and lung inflammation, such as Integrin and CCKR (cholecystokinin receptor) signaling. More interestingly, using single-cell transcriptomics data from the human cell atlas, the EV-specific transcriptomic signature of pre-RESP samples distinctly associated with chronic obstructive pulmonary disease and corresponded to systemic activation of cell types of the innate immune system, revealing disease-, cell-, and tissue-specificity of the underlying systemic signaling and pathophysiological response. The findings highlight the potential of urinary EVs to capture dynamic changes in pathophysiological responses, including the underlying cell-to-cell signaling, in chemical inhalational exposures. Expansion of this proof-of-concept to other clinical and subclinical conditions can lead to development of transformative EV-based liquid biopsies for longitudinal non-invasive biomonitoring of the underlying organismal signaling in exposures, diseases, and therapeutic treatments, enabling highly personalized interventions and therapeutic strategies across chemical toxicology and health.

## RESULTS

### Study participants and exposures

The demographics of study participants are summarized in **Table 1**. The five workers, all male, were self-identified as white of Hispanic/Latino origin with an average group age of 41 years (range 30-53 years). These workers were on the job for an average of 2.5 years (range 1 to 6 years). Inhalation exposures of these workers to MDI were low (**Table 1**), with a geometric mean (GM) ± geometric standard deviation (GSD) of 18.65 ± 6.6 ng/m^3^. These airborne exposures translate to ∼2 parts per trillion (2 ppt) airborne MDI.

**Table 1.** Demographics and summary of inhalation exposures of chronically exposed workers in the study

|  |  | Chronically exposed workers |
| --- | --- | --- |
| **Age (years)** | Mean ± SD | 41.2 ± 3.9 |
| **Gender (#)** | Male | 5 |
|  | Female | - |
| **Race (#)** | White | 5 |
|  | Asian | - |
|  | Black | - |
| **Ethnicity (#)** | Hispanic or Latino | 5 |
|  | Non-Hispanic or Latino | - |
| **Current smokers (#)** |  | - |
| **Years on job** | Mean ± SD | 2.5 ± 2.5 |
| **Inhalation MDI exposures** | PBZ inhalation (ng/m<sup>3</sup>)<br>(Geometric Mean ± SD) | 18.65 ± 6.6 |

MDI – methylene diphenyl diisocyanate; PBZ – personal breathing zone

### Biophysical characterization of extracellular vesicles

The overall study design for respirator intervention and urine sample collection is shown in **Fig. 1A**. Purified urinary EVs were characterized as per the guidelines of the International Society of Extracellular Vesicles (ISEV) (*23*). EV size and morphology were evaluated using transmission electron microscopy (TEM) and cryo-TEM and were confirmed to represent vesicle-shaped structures in the 50 – 200 nm size range with some vesicles being multilamellar (**Fig. 1B-C**). The urinary EVs were found to be significantly enriched in established EV markers (CD63, CD9, and CD81) and depleted of cytosolic protein, GAPDH, further confirming the purity of the isolated EVs (**Fig. 1D**). The size distribution of the purified EVs observed in TEM was also confirmed by TRPS (tunable resistive pulse sensing) analysis (*24*). In the TRPS analysis, the average size distribution and concentration were very similar between pre-RESP and post-RESP samples, although there was higher within-sample variability in EV concentration of pre-RESP samples (**Fig. 1E**, left panel). However, the average total concentration of EVs was 6.65 (± 3.1) × 10^10^ per mL for pre-RESP and 1.35 (± 0.63) × 10^11^ per mL for post-RESP samples, which was not significantly different (**Fig. 1E**, right panel). We also conducted creatinine content analysis of the urine samples as per the recommendation of the Urine Task Force of the ISEV (*25*, *26*), and is summarized in **Table S1**. Lastly, cellular component ontology analysis of all identified EV-proteins in proteomics analysis indicated statistically robust enrichment for extracellular vesicles (p-value = 5e-324) (**Fig. 1F** and bioinformatics supplementary data), indicating high purity of urinary EV isolations.

**Figure 1.**
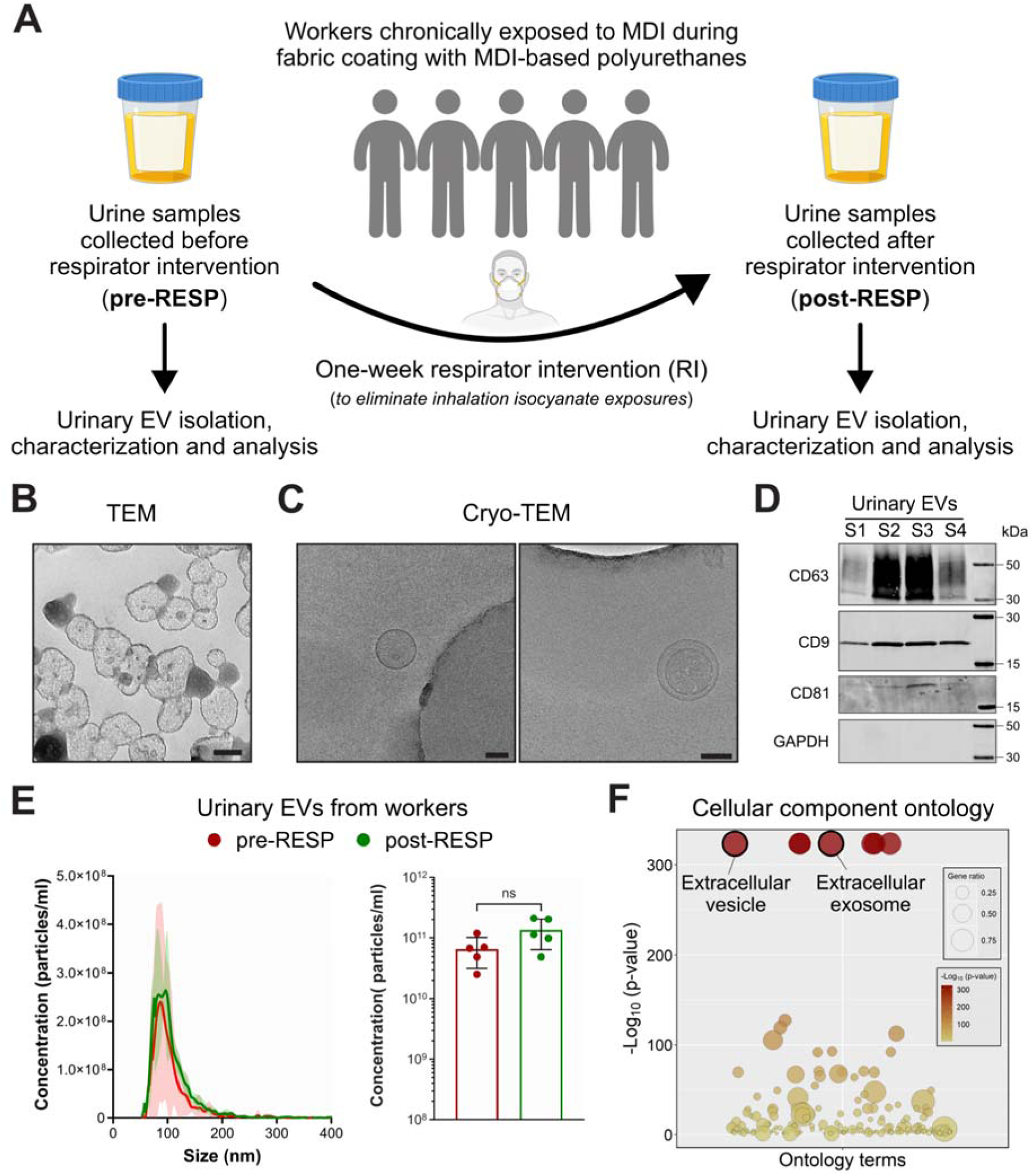
Biophysical characterization of urinary extracellular vesicles. (**A**) Overall schematic of the study design. MDI – methylene diphenyl isocyanate. (**B**) Representative transmission electron microscope (TEM) and (**C**) cryo-TEM images of purified urinary EVs (size bar - 100 nm). (**D**) Representative western blot of urinary EVs from four different subjects demonstrating the presence of EV markers (CD63, CD9, and CD81) and absence of cytosolic marker, GAPDH. Twenty percent of the total EV pellet was used for evaluation. (**E**) Average size distribution and concentration of purified urinary EVs (left panel) as well as total particle concentration (right panel) from pre-RESP and post-RESP samples (n = 5 per group). Error bars: mean ± SD, Welch’s two-tailed t-test. (**F**) Cellular component ontology analysis of all identified EV-proteins indicates very high purity of the EV isolations.

### EV-mRNAs differentially abundant in pre-RESP samples are associated with inflammatory airway disease-related biological processes

We performed next generation deep sequencing of total RNA from EVs of pre-RESP and post-RESP urine samples. The overall percentage distribution of unique reads was similar across all samples (**Fig. S1A-B**). We first focused on the mRNA content of the EVs. While the principal component analysis (PCA) of EV-mRNAs showed only moderate separation between the two groups (**Fig. 2A**), we focused on EV-mRNAs that were significantly differentially abundant in the pre-RESP vs post-RESP comparison (**Fig. 2B**). Globally, 393 EV-mRNAs were significantly enriched and 372 EV-mRNAs were significantly depleted in the pre-RESP samples compared to the post-RESP samples (**Fig. 2B-C**). Some of the top EV-mRNAs enriched or depleted in pre-RESP samples have already been implicated in inflammatory airway disease. AP3B1, which was the most enriched mRNA in pre-RESP EV samples (**Fig. 2D**), has been strongly linked to chronic obstructive pulmonary disease (COPD) risk in the largest genome-wide association study of lung function to date (*27*). USP7, which was highly enriched in pre-RESP samples, has been reported to be upregulated in COPD tissues and inhibiting USP7 alleviated airway mucus hypersecretion in *in vitro* and *in vivo* models (*28*). Similarly, EV-mRNAs significantly depleted in pre-RESP samples, such as HSF1 and FFAR4, have been reported to play a protective role in inflammatory airway diseases (*29–31*). Consistent with these findings, panther pathway analysis of the differentially enriched EV-mRNAs demonstrated significant enrichment for several pathways implicated in inflammatory airway diseases, such as integrin signaling, hedgehog signaling, endothelin signaling and CCKR signaling pathways (**Fig. 2E**). Collectively, the results indicate marked changes in inflammatory airway disease-associated mRNAs in the urinary EVs following respiratory exposure reduction. It further confirms sufficient sensitivity of the urinary EV-mRNA analysis for the purpose of documenting changes in biological responses following exposure interventions to minimize inhalation exposures to MDI and other airborne toxicants.

**Figure 2.**
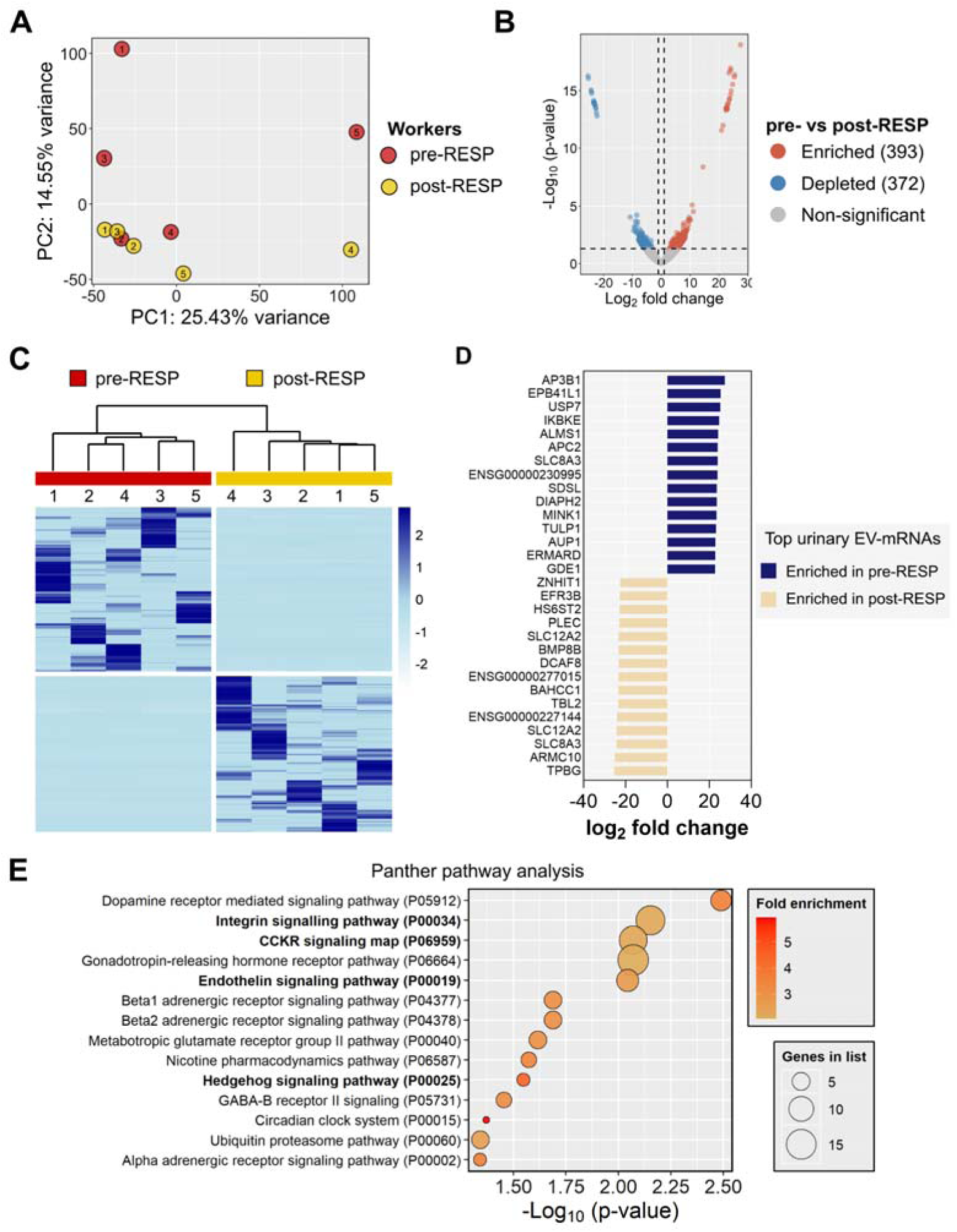
EV-mRNA abundance markedly changes following respirator intervention and correlates with inflammatory airway disease-related pathways. (**A**) Principal Component Analysis using top 2500 variable EV-mRNAs (2.5%) demonstrates modest clustering of post-RESP samples and outgrouping of three pre-RESP samples. (**B**) Volcano plot of significantly differentially enriched mRNA transcripts between pre-RESP and post-RESP samples. The transcripts are plotted according to their -log_10_(p-values) as determined by two-tailed t-test and log_2_ fold change. Red dots: mRNA transcripts significantly enriched identified with p >0.05 and log_2_ fold change > 1. Blue dots: mRNA transcripts significantly depleted identified with p >0.05and log_2_ fold change < -1. Grey dots: mRNA transcripts not significantly differentially abundant. (**C**) Hierarchical clustering and heatmap using significantly differentially enriched mRNA transcripts in pre-RESP vs post-RESP comparison (p < 0.05, distance method = manhattan, clustering method = ward.D2). (**D**) A curated list of top 15 EV-mRNAs enriched in pre-RESP or post-RESP samples. (**E**) Overrepresentation analysis based on significantly differentially enriched EV-mRNAs identifies panther pathways statistically significant and overrepresented in pre-RESP vs. post-RESP comparison. The pathways are plotted based on their p-value (Fisher’s t-test with no FDR correction). Fold enrichment and number of genes identified per Panther pathway are indicated.

### EV-mRNAs distinctly enriched in pre-RESP group reveal global immune response mediated by monocytes, macrophages, and a subset of neutrophils

The integrated human lung cell atlas includes transcriptomics data of healthy lung tissue and of various lung diseases. So, we asked if urinary EV-based transcriptome signature of the workers in this study demonstrates specificity to inflammatory airway-related diseases and does respirator intervention changes that signature. To evaluate this question, we selected the top 50 EV-mRNAs enriched in the pre-RESP (pre-RESP^high^ EV-mRNAs) or post-RESP (post-RESP^high^ EV-mRNAs) samples. Indeed, pre-RESP^high^ EV-mRNAs, but not post-RESP^high^ EV-mRNAs, were distinctly overexpressed in chronic obstructive pulmonary disease (**Fig. 3A**). Since the lung cell atlas contains single-cell and spatial transcriptomics data (**Fig. 3B**), we then asked whether EV-mRNA signature can reveal cellular and tissue specificity of an underlying pathophysiological response. Interestingly, we found that pre-RESP^high^ EV-mRNAs, but not post-RESP^high^ EV-mRNAs, were overexpressed primarily in myeloid leukocytes, innate lymphoid cells, and granulocytes (**Fig. 3C-D** and **Fig. S2A**). On the other hand, the post-RESP^high^, but not pre-RESP^high^, EV-mRNAs were found to overexpressed primarily in adventitial cells of the lung (**Fig. 3D** and **Fig. S2A**). Since urine and the derived EVs can also be highly reflective of kidney health, we also conducted the analysis using the kidney cell atlas (**Fig. 3E**). No distinct differences were observed in kidney-resident epithelial and endothelial cells (**Fig. S2B**). However, similar to the lung cell atlas, the myeloid leukocytes and innate lymphoid cells were found to overexpress pre-RESP^high^ EV-mRNAs (**Fig. 3F** and **Fig. S2B**), which led us to hypothesize that the pre-RESP-specific EV-mRNA signature is indicative of a global immune response to inhaled MDI and other co-toxicants.

**Figure 3.**
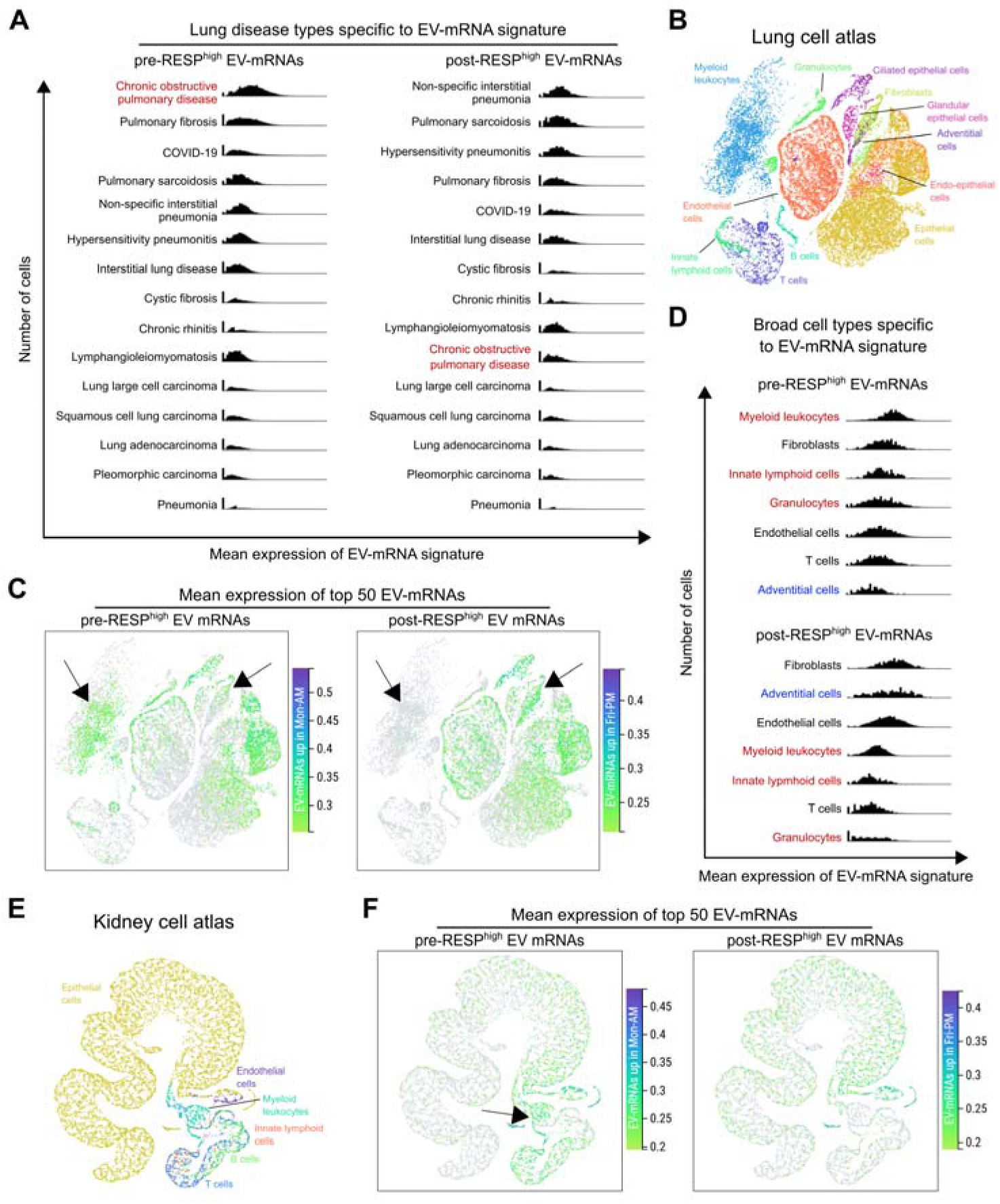
EV-mRNAs highly enriched in pre-RESP samples largely correspond to cell types of the innate immune system in lung and kidney cell atlases. (**A**) UMAP-based visualization and clustering of single-cell expression data in human lung cell atlas. Individual dots represent each cell and the colors of the dots represent the broad classes to which the cells belong. (**B**) Mean expression of top 50 pre-RESP- (pre-RESP^high^) or post-RESP-enriched (post-RESP^high^) EV-mRNAs in various cell classes in the lung cell atlas. Arrows indicate differential enrichment of pre-RESP^high^ or post-RESP^high^ EV-mRNAs among the various cell classes. Only cells in the top 10 percentile based on mean EV-mRNA expression are shown as colored dots. (**C**) Histogram-based representation of the mean expression of pre-RESP^high^ or post-RESP^high^ EV-mRNAs in various classes of cells in the lung, sorted by highest to lowest mean expression. Cell classes highlighted in red indicate higher expression of pre-RESP^high^ EV-mRNAs and in blue indicate higher expression of post-RESP^high^ EV-mRNAs. (**D**) UMAP-based visualization and clustering of single-cell expression data in human kidney cell atlas. Individual dots represent each cell and the colors of the dots represent the broad class to which the cells belong. (**E**) Mean expression of pre-RESP^high^ or post-RESP^high^ EV-mRNAs in various cell classes in the kidney cell atlas. Arrow indicates differential enrichment of pre-RESP^high^ or post-RESP^high^ EV-mRNAs among myeloid leukocytes. Only cells in the top 10 percentile based on mean EV-mRNA expression are shown as colored dots. (**F**) Histogram-based representation of the mean expression of pre-RESP^high^ EV-mRNAs in various classes of cells in the kidney, sorted by highest to lowest mean expression. Cell classes highlighted in red indicate higher expression of pre-RESP^high^ EV-mRNAs.

To further substantiate this observation, we evaluated the expression of pre-RESP^high^ EV-mRNAs and post-RESP^high^ EV-mRNAs in the global immune cell atlas (**Fig. 4A**). While post-RESP^high^ EV-mRNAs were not enriched in any immune cell type, pre-RESP^high^ EV-mRNAs were found to be highly overexpressed in specific immune cell types (**Fig. 4B**). These included myeloid leukocytes, hematopoietic cells, and a subset of granulocytes (**Fig. S2C**). To evaluate our hypothesis of a global immune response, we determined whether the enrichment of pre-RESP^high^ EV-mRNAs in these cell types was global or tissue-specific. Among myeloid leukocytes, the pre-RESP-specific EV-mRNA signature was found to be high in classical monocytes in the blood and lung; intermediate monocytes in the lung; tissue-resident macrophages in parotid gland; monocytes in blood, lung, spleen, liver, and coronary artery; mononuclear phagocytes in the stomach; and macrophages in the blood, lung, exocrine pancreas, and coronary artery (**Fig. 4C-E** and **Fig. S3**). In the same cell types, post-RESP^high^ EV-mRNAs were not observed to be highly expressed (**Fig. S4A**). In hematopoietic cells, hematopoietic precursor cells exhibited distinctly high expression of the pre-RESP^high^ EV-mRNA, but not post-RESP^high^ EV-mRNA signature, primarily in the bone marrow (**Fig. 4F** and **Fig. S4B**). And lastly, in granulocytes, the pre-RESP-specific EV-mRNA signature was high in primarily neutrophils in the bone marrow (**Fig. 4G** and **Fig. S4B**). In conclusion, the pre-RESP^high^ EV-mRNA signature distinctly associated with chronic obstructive pulmonary disease and corresponded to cell types of the innate immune system with global expression in monocytes and macrophages across several tissues and neutrophils and hematopoietic precursor cells in the bone marrow, suggesting systemic activation of the hematopoietic system in response to an immune response.

**Figure 4.**
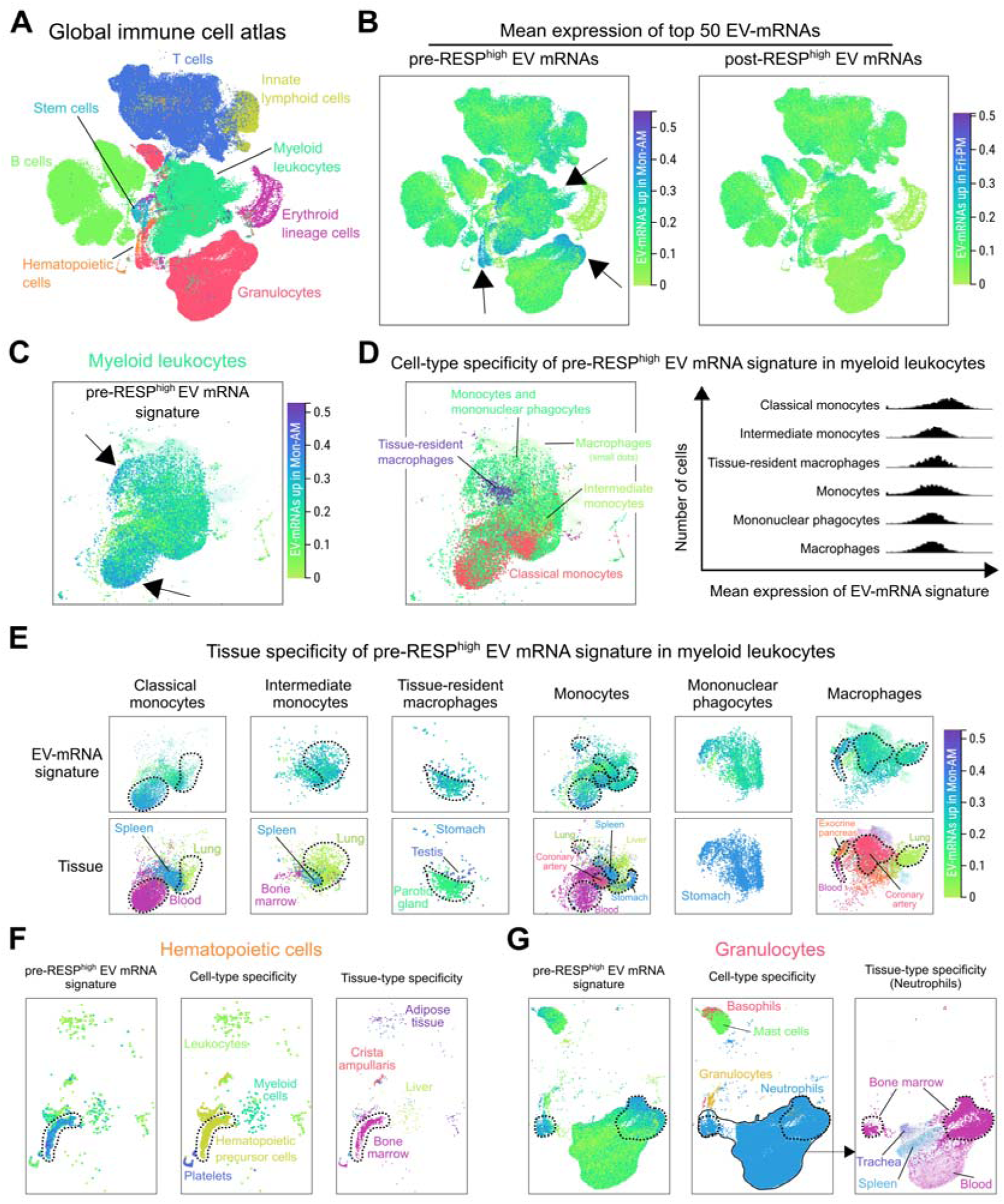
Correlation of pre-RESP^high^ EV-mRNAs with immune cell atlas suggests global immune response in workers before mask intervention. (**A**) UMAP-based visualization and clustering of single-cell expression data in global immune cell atlas. Individual dots represent each cell and the colors of the dots represent the broad classes to which the cells belong. (**B**) Mean expression of pre-RESP^high^ or post-RESP^high^ EV-mRNAs in various cell classes in the immune cell atlas. Arrows indicate differential enrichment of pre-RESP^high^ or post-RESP^high^ EV-mRNAs among myeloid leukocytes, hematopoietic cells, and granulocytes. (**C**) An expanded view of the mean expression of pre-RESP^high^ EV-mRNAs in myeloid leukocytes and (**D**) the specific cell types among the myeloid leukocytes class that demonstrate high expression of pre-RESP^high^ EV-mRNAs, i.e. classical monocytes, intermediate monocytes, tissue-resident macrophages, monocytes, mononuclear phagocytes, and macrophages. In the histogram-based representation, the cell types are sorted by highest to lowest mean expression. (**E**) Mean expression of pre-RESP^high^ EV-mRNAs in myeloid leukocytes described in D, and its correlation with tissue-based UMAP to determine specific tissues where pre-RESP^high^ EV-mRNA signature is high for each cell type. (**F** and **G**) Correlation between mean pre-RESP^high^ EV-mRNA expression, cell-type-based UMAP, and tissue-based UMAP of (**F**) hematopoietic cells and (**G**) granulocytes to determine spatial specificity of pre-RESP^high^ EV-mRNA signature.

### Identification of differentially abundant non-coding RNAs

We then focused on the non-coding RNA (ncRNA) content of the urinary EVs before and after respiratory exposure reduction (pre-RESP and post-RESP), which included microRNAs (miRNAs), long intergenic non-coding RNAs (lincRNAs), small nuclear RNAs (snRNAs), small nucleolar RNAs (snoRNAs), ribosomal RNAs (rRNAs), antisense RNAs (asRNAs) and other RNAs. Similar to mRNA, principal component analysis only moderately separated the two groups (**Fig. 5A**), however, a significant number of non-coding RNAs were found to be statistically differentially enriched between the pre-RESP and post-RESP samples (**Fig. 5B**). Of the total significantly differentially enriched ncRNAs (163), 66% (107) were depleted and 34% (56) were enriched in the pre-RESP samples in comparison to post-RESP samples (**Fig. 5B-D**). While it remains unclear if the differentially enriched ncRNAs are correlated or associated with inhalation exposures, mainly due to lack of a comprehensive understanding of the role of ncRNAs in pathophysiological processes and lack of ncRNA-based ontology analysis tools, existing literature correlated with our findings. SNHG5 – a snoRNA, which was one of the most significantly depleted ncRNAs in pre-RESP samples, has been reported to be downregulated in chronic obstructive pulmonary disease (COPD) patient tissue and serum samples, and has been proposed as a novel prognostic biomarker for acute exacerbation of COPD (*36*, *37*). On the other hand, PLCE1-AS2 – an asRNA, was one of the most significantly enriched ncRNAs in pre-RESP samples. While it remains unclear how PLCE1-AS2 regulates PLCE1 transcription or translation, the expression of PLCE1 is upregulated in acute lung injury model and its suppression alleviates acute lung injury through inhibition of PKC and NF-κB signaling pathways (*38*). These results suggest that in addition to mRNA, the ncRNA content of urinary EVs may reflect changes in inhalation exposures of individuals and they may provide additional information on the potential physiological responses and immune system remodeling that follows changes in inhalation exposures.

**Figure 5.**
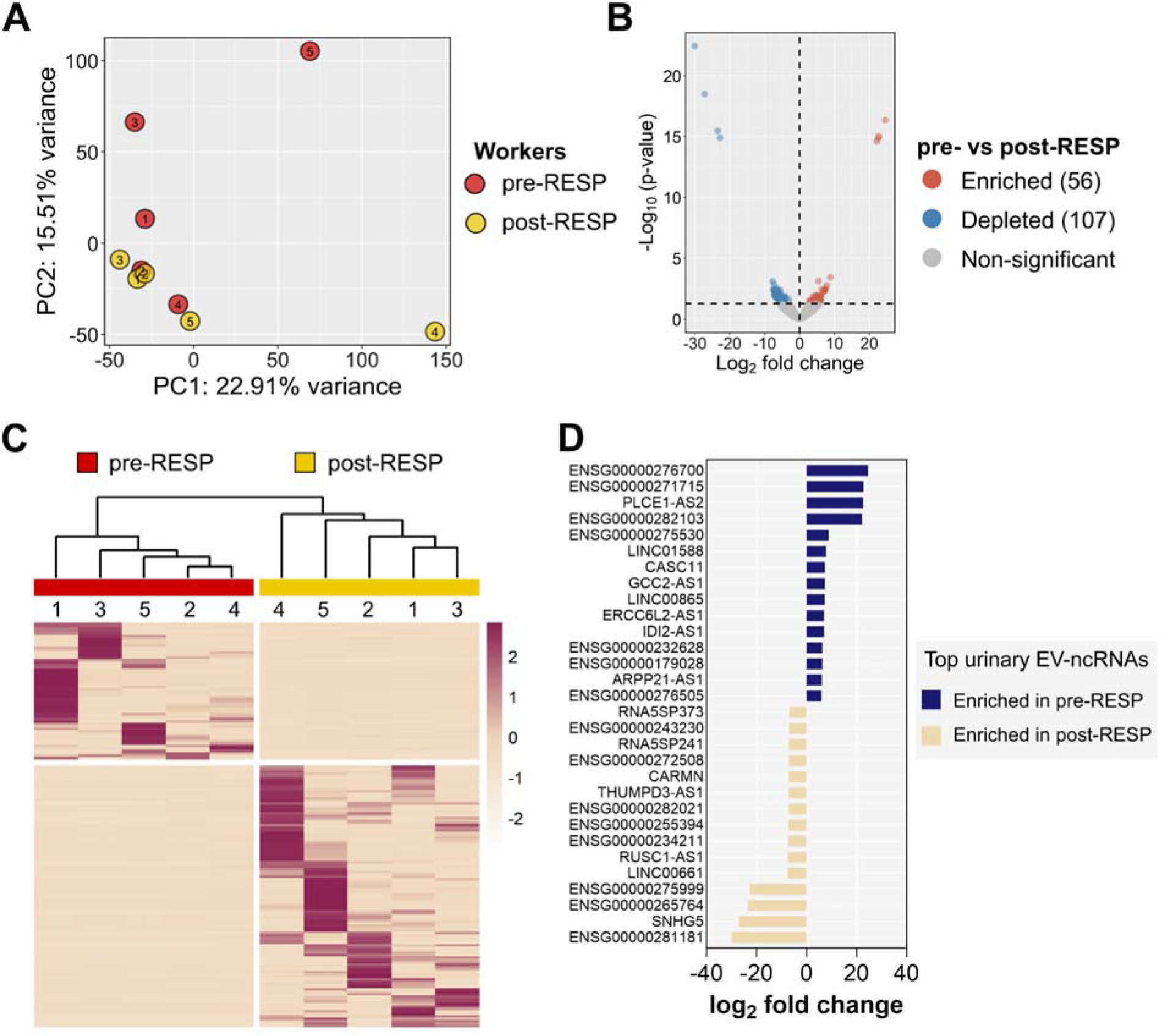
Abundance of EV-ncRNAs remarkably changes following respirator intervention. (**A**) Principal Component Analysis using all EV-specific non-coding RNAs (ncRNAs) demonstrates modest clustering of post-RESP samples and outgrouping of three pre-RESP samples. (**B**) Volcano plot of significantly differentially enriched ncRNAs between pre-RESP and post-RESP samples. The transcripts are plotted according to their -log_10_(p-values) as determined by two-tailed t-test and log_2_ fold change. Red dots: ncRNAs significantly enriched identified with p >0.05and log_2_ fold change > 1. Blue dots: ncRNAs significantly depleted identified with p >0.05and log_2_ fold change < -1. Grey dots: ncRNAs not significantly differentially abundant. (**C**) Hierarchical clustering and heatmap using significantly differentially enriched ncRNAs in pre-RESP vs post-RESP comparison (p < 0.05, distance method = manhattan, clustering method = ward.D2). (**D**) A curated list of top 15 EV-ncRNAs enriched in pre-RESP or post-RESP samples.

### Distinct EV proteome of pre-RESP samples also associated with inflammatory airway disease-related biological processes

Lastly, we evaluated if changes in the proteome of the pre-RESP and post-RESP urinary EV samples correlated with changes in isocyanate chemical exposure. In comparison to PCA of EV-mRNAs and -ncRNAs, proteome-based PCA led to a more distinct separation between the pre-RESP and post-RESP groups (**Fig. 6A**). A total of 74 proteins were found to be statistically differentially enriched between the two groups, of which 27 were enriched and 47 were depleted in the pre-RESP samples compared to the post-RESP samples (**Fig. 6B-C**). MT1G, the most enriched EV-protein in pre-RESP samples, and other members of the MT1 family have been suggested to have a broader role in inflammatory diseases and shaping innate and adaptive immune responses (*39*). RNASE7, another highly abundant protein in pre-RESP samples, has also been reported to be overexpressed in basal cells of airway epithelia in response to damaged airway epithelium (*40*). Another pre-RESP-enriched EV-protein, EPPK1, is upregulated in normal bronchial epithelial cells upon exposure to cigarette smoke (*41*). One the other hand, SLURP1, which was the most enriched EV-protein in post-RESP samples has been shown to inhibit the progression of inflammation by influencing both neutrophils and endothelial cells at multiple steps, ultimately resulting in suppression of neutrophil extravasation and chemotaxis (*42*). SERPINB6, another post-RESP-enriched EV-protein, has also been suggested to be protective against inflammation through its ability to inhibit peptidases stored in cytolytic granules in monocytes and granulocytes (*43*, *44*). Panther pathway analysis of the differentially enriched EV-proteome demonstrated significant enrichment for inflammatory airway disease-related pathways that were also identified in the EV-mRNA analysis. This included integrin signaling and CCKR signaling pathways (**Fig. 6D**). Lastly, we determined if the corresponding mRNAs of EV-proteins enriched in pre-RESP or post-RESP samples demonstrate cell-specific enrichment in the human lung cell atlas. Interestingly, the corresponding transcripts for pre-RESP-enriched EV-proteins were distinctly enriched in myeloid leukocytes and in granulocytes to some extent (**Fig. 6E** and **Fig. S5**). In contrast, the transcripts for post-RESP-enriched EV protein signature were highly enriched in adventitial cells and fibroblasts. These findings demonstrate that the urinary EV proteome of the workers markedly changes within a week following respirator intervention and provides both overlapping and distinct insights when compared to EV-mRNA analysis.

**Figure 6.**
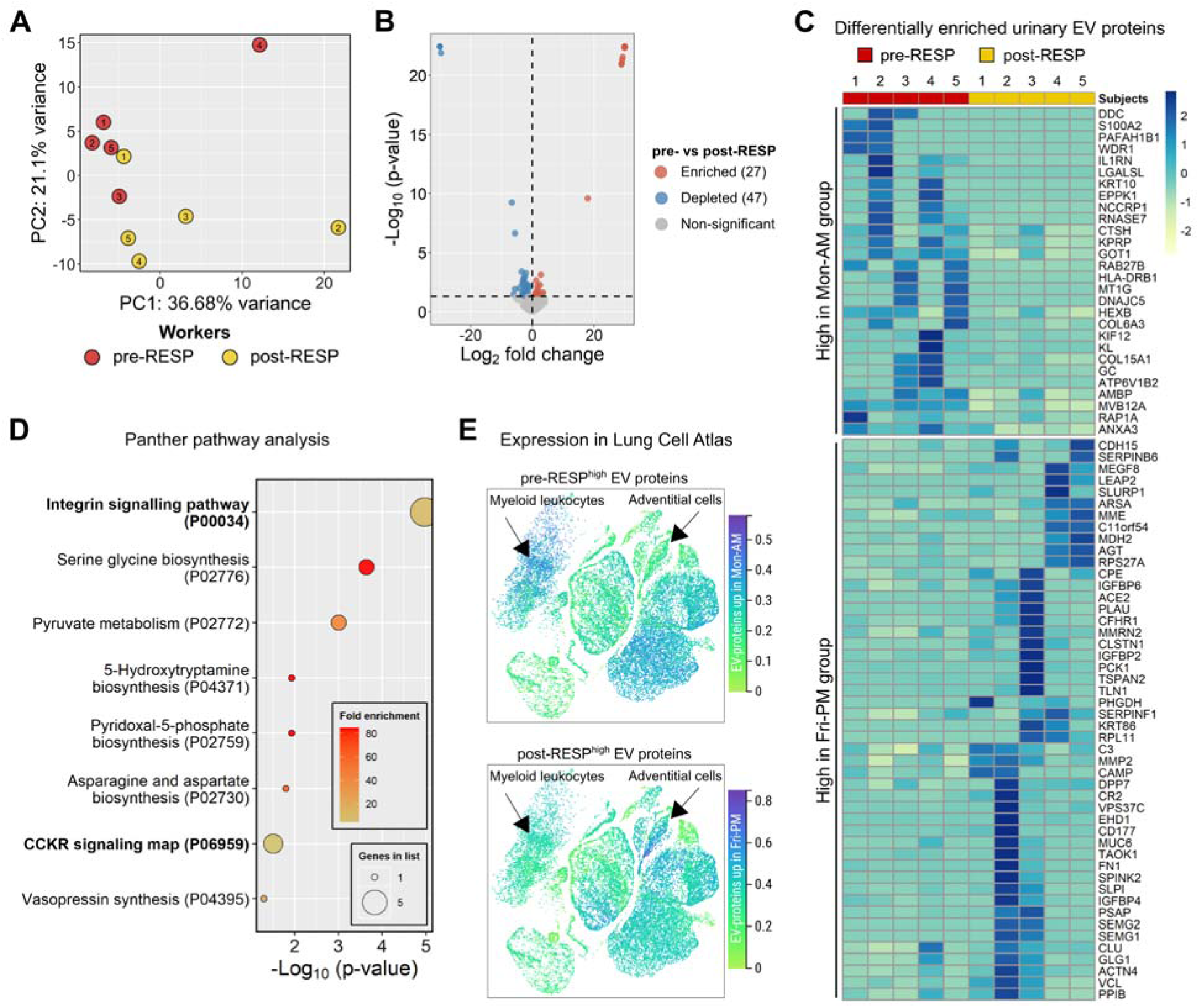
EV-proteome also undergoes significant changes following respirator intervention and is associated with inflammatory airway disease-related pathways. (**A**) Principal Component Analysis using all EV-specific proteins demonstrates modest separation between pre-RESP and post-RESP groups. (**B**) Volcano plot of significantly differentially enriched proteins between pre-RESP and post-RESP samples. The proteins are plotted according to their -log_10_(p-values) as determined by two-tailed t-test and log_2_ fold change. Red dots: proteins significantly enriched identified with p >0.05 and log_2_ fold change > 0. Blue dots: proteins significantly depleted identified with p >0.05 and log_2_ fold change < 0. Grey dots: proteins not significantly differentially abundant. (**C**) Hierarchical clustering and heatmap using significantly differentially enriched proteins in pre-RESP vs post-RESP comparison (p < 0.05, distance method = manhattan, clustering method = ward.D2). (**D**) Overrepresentation analysis based on significantly differentially enriched EV-proteins identifies panther pathways statistically significant and overrepresented in pre-RESP vs. post-RESP comparison. The pathways are plotted based on their p-value (Fisher’s t-test with no FDR correction). Fold enrichment and number of genes identified per Panther pathway are indicated. (**E**) Mean expression of pre-RESP^high^ EV-proteins (27) or post-RESP^high^ EV-proteins (47) in various cell classes in the lung cell atlas. Arrows indicate the cell classes that demonstrate differential enrichment of pre-RESP^high^ or post-RESP^high^ EV-proteins.

## DISCUSSION

While EVs are being evaluated as prognostic and diagnostic biomarkers of established clinical diseases, utilization of EVs and their cargo to probe biological responses and remodeling of immune system communication pathways following inhalational exposures remains understudied. In this pilot work, we optimized methodologies for urinary EV isolation as well as analysis of EV-derived RNAs and proteome in five workers with well-characterized chronic exposures to inhaled toxicants in an MDI fabric coating factory. The urine samples were collected from the same individuals before (pre-RESP) and after (post-RESP) a one-week of OVC respirator intervention to minimize inhalation exposures, allowing us to uniquely evaluate temporal changes in the urinary EV cargo following inhalation exposure reduction due to intervention. The application on isocyanate-exposed workers is particularly of interest considering isocyanates are potent sensitizers and there are no easy tests and reliable early biomarkers for detection of sensitized individuals. Even more interesting in this exposure scenario is extremely low inhalation exposures to MDI (∼2 ppt).

Our results consistently indicated distinct enrichment of EV-mRNAs, -ncRNAs, and -proteins within one week of respirator intervention to reduce inhalation exposures. Furthermore, the analyses of differentially enriched EV cargoes exhibited strong correlations with several biomolecules reported to be directly involved in lung disease like COPD in *in vitro*, *in vivo*, and patient studies. In longitudinal studies of healthy individuals, urinary EV proteome has been reported to be stable over several months (*45*, *46*), suggesting that the cross-week changes that we observe in the urinary EV cargo are associated with activation of homeostatic biological responses during this one week inhalation exposure reduction intervention, which can be captured non-invasively through urinary EV analysis.

Our study clearly demonstrates high sensitivity of the urinary EV cargo as seen in the cross-week changes in urinary EV-mRNA, -ncRNA, and -proteome upon respiratory exposure intervention in chronically exposed individuals. Additionally, given the role of EVs in intercellular communication, a major goal in the EV field has been to use EV-based omics to delineate the underlying signaling, and the cell types involved. Its realization, however, has remained challenging due to various fundamental reasons related to EV heterogeneity and other factors. Recent advancements in single-cell transcriptomics, including the publication of the first draft of the human cell atlas – a comprehensive reference map of all cells – has created unique opportunities in diagnosing, treating, and monitoring diseases. In this study, we propose an approach that pairs EV-specific transcriptomes with the human cell atlas to reveal the identity of the underlying intercellular signaling. The human cell atlas represents 18 single-cell-transcriptome-based maps of different important organs and tissues in healthy and diseased states. By pairing EV-specific transcriptomes with the human cell atlas, we demonstrated that urinary EV-mRNAs specifically enriched in chronically exposed worker samples (pre-RESP) are strongly associated with the mRNA signature of COPD. Moreover, the pre-RESP-specific EV transcriptome was highly expressed in cell types of the innate immune system. These included monocytes and macrophages across several tissues and neutrophils and hematopoietic precursor cells in the bone marrow. Indeed, chronic exposure to isocyanates and its albumin adducts are well documented to lead to a hypersensitive state of the immune system, characterized by marked changes in monocyte and macrophage function (*47*, *48*), neutrophil exhaustion and neutrophilia (*49*, *50*), and activation of B and T cell responses (*51–53*). Future experimental validation of this approach can provide novel ways for constant non-invasive biomonitoring of exposures, diseases and therapeutic responses, and thereafter, providing personalized interventions and therapies.

While the findings are promising, the study has some inherent limitations, and the findings should be interpreted with caution and only in the context of methodological developments. First, although the small size of the cohort was intentional due to the pilot nature of the study, significant variation in their chronic exposures ranging from 1 to 6 years could have influenced our analyses. Future studies in larger cohorts will provide a comprehensive and reliable evaluation of urinary EVs for biomarker discovery and biomonitoring purposes in chemical toxicology. Second, in addition to reducing inhalation exposures to isocyanates, the respirator intervention could have also reduced exposures to other chemicals and particulate matter in the factory that were not measured as part of the study. Therefore, the changes in urinary EV cargo that we observe are likely reflective of overall reduction in inhalation exposures. Third, while the respirator intervention significantly reduced inhalation exposure, the study design did not control for skin exposure during the same time, which could influence our urinary EV cargo analysis. Lastly, complementing urinary EVs with inflammatory markers in circulation and urinary kidney injury biomarkers will provide more clinically-relevant context to the observed RNA and proteome changes. Ultimately, we think urinary EVs and their cargo may pave the way for the discovery and development of non-invasive screening tests for early detection of isocyanate exposed and/or sensitized workers at high risk of developing occupational asthma.

In conclusion, our study demonstrates the feasibility and sensitivity of urinary EVs and their contents (mRNAs, ncRNAs and proteins) for capturing short-term (such as cross-week) changes in underlying systemic signaling following inhalation exposure reduction and their promise for biomarker discovery and applications in chemical toxicology. Expansion of this proof-of-concept paradigm that links urinary EV-based transcriptome, proteome, and later metabolome, with a targeted exposome could provide a framework for a deeper and more comprehensive analysis of systemic biological responses to toxins using urine, a noninvasive biological medium, which in certain settings may be the only medium that can be collected.

## MATERIALS AND METHODS

### Subject recruitment, exposure characterization and urine collection

Study participants were recruited and signed an informed consent approved by the Institutional Review Board at Yale University (protocol number 0311026133) and University of Massachusetts Lowell, as previously described (*22*). Daily personal inhalation exposure to MDI, workplace description, and routine medical evaluation of blood histochemistry for MDI-specific antibody testing, and pulmonary and liver function are detailed in the previously published studies (*22*, *54*). Spot urine samples from each worker were collected in sterile urine specimen collection cups as previously described. For this study, urine samples from five workers collected before and after respirator intervention.

### Creatinine analysis

The protocol for the LC-ESI-MS/MS of creatinine was based on previously published literature (*55*). Briefly, the thawed urine samples were diluted first by 100× (10 µL into 990 µL DI water), followed by a second 20× dilution (50 µL into 1 mL final) in LC amber glass vials. Ten nanogram of creatine-d3 were spiked into this second LC vial (final concentration, 10 ng/mL). This urine solution was subject to LC-ESI-MS/MS analysis (*55*).

### Isolation of extracellular vesicles from urine

Cell-free urine samples stored at -80°C were thawed on ice. EVs were isolated using a qEVoriginal (70nm) size exclusion column (Izon Science Ltd.). After washing the qEV column with 1X PBS, 500 μl of thawed urine sample was added to the column. After discarding the first 3 ml of flow through (equivalent to six fractions), considered as void volume, EV fraction of 1,500 μl was collected. Isolated EVs were characterized for morphology, size distribution, and concentration as described in ‘EV characterization’ section.

For RNA isolation and western blotting analysis, 5 ml of thawed urine sample was transferred to an ultracentrifuge tube (Beckman Coulter, Inc.). The sample was diluted with 25 ml of nuclease-free 1X PBS (Thermo Fisher Scientific, AM9624) and filtered through a 0.2 µm filter. The diluted urine samples were centrifuged at 200,000 ×g for 65 min at 4°C to pellet EVs. The pellet was resuspended in 250 μl or 150 μl of nuclease free water for RNA isolation or 1X PBS for wester blotting.

### Extracellular vesicle characterization

Morphological analysis of EVs was conducted using transmission and cryo-electron microscopy, whereas size distribution and concentration were determined using qNano instrument (Izon Science Ltd.). For transmission electron microscopy (TEM), 5 μl of purified urinary EVs from above were added onto carbon-coated 400-mesh copper grids (Electron Microscopy Sciences, CF400-Cu) and air-dried. The grids were then visualized using Philips CM12 transmission electron microscope at 100 keV. Cryo-electron microscopy analysis of isolated EVs was performed at the Harvard Center for Nanoscale Systems. For cryo-electron microscopy, the purified EVs were applied on a glow-discharged (2 minutes in 0.2 mbar air using a EMITECH K950X with glow discharger unit) carbon-coated 400-mesh copper grids (Electron Microscopy Sciences, CF400-Cu) and were vitrified using a Gatan CP3 cryoplunger (Gatan, Inc.) at room temperature and 100% humidity. Excess sample was removed by blotting once between 1 and 2 seconds with filter paper. The blotted grids were plunged into liquid ethane that was kept in equilibrium with solid ethane. After vitrification, the grid was stored under liquid nitrogen until further use. Grids were visualized using a FEI Tecnai Arctica CryoTEM (Thermo Fisher Scientific, Inc.) with Autoloader at 200 keV.

The size distribution of purified EVs was further characterized using a qNano instrument (Izon Science Ltd.) fitted with a NP100 nanopore (size range of 50-200 nm). Polystyrene beads of 100 nm size (Thermo Fisher Scientific, 3100A) were used before and after sample for calibration and EV number concentration (# of EVs/ml), and 0.2 μm-filtered 1X PBS was used as a negative control.

EVs were also evaluated for the presence of EV-enriched markers (CD63, CD9, and CD81) and the absence of EV-excluded proteins (GAPDH). Blots were incubated in respective antibodies and imaged using Odyssey^®^ CLx instrument and Image Studio software (LI-COR, Inc.). Images were acquired with blots facing down on scan bed and the following scan control settings – scan resolution ‘169 µm’; scan quality ‘high’; and focus offset ‘0.0 mm’. Cut, uncropped blots, depicting the full region scanned on the Licor can be found in Supplementary Information. The following antibodies were used – anti-CD9 rabbit monoclonal antibody (Cell Signaling, cat no. 13174), anti-CD81 mouse monoclonal antibody (Thermo Fisher Scientific, cat no. 10630D), anti-CD63 mouse monoclonal antibody (Fisher Scientific, cat no. NBP232830F), and anti-GAPDH rabbit monoclonal antibody (Cell Signaling, cat no. 2118S) as primary antibodies; and IRDye^®^ 680RD Goat anti-Mouse IgG (LI-COR, cat no. 926-68070), IRDye® 680RD Goat anti-Rabbit IgG (LI-COR, cat no. 926-68071), IRDye^®^ 800CW Goat anti-rabbit IgG (LI-COR, cat no. 926-32211), and IRDye® 800CW Goat anti-Mouse IgG (LI-COR, cat no. 926-32210) as secondary antibodies.

### EV cargo analysis of RNAs using next generation deep sequencing

#### Total RNA isolation and library preparation

Total RNA from EV pellets was isolated using MagMAX™ mirVana™ Total RNA Isolation Kit (Thermo Fisher Scientific, A27828) and eluted in 10 μL of elution buffer. After confirming that small RNAs were abundant in the isolated urinary EV total RNA by using Agilent RNA 6000 Nano Kit (Agilent Technologies, 5067-1511), the concentration and size distribution of isolated RNA from all samples was quantified with an Agilent BioAnalyzer system with a small RNA kit (Agilent Technologies, 5067-1548). To prevent exclusion of RNAs lacking 5’phosphate or 3’hydroxyl, purified RNA was treated with T4 polynucleotide kinase (PNK) according to manufacturer’s instructions (Thermo Fisher Scientific, EK0031), further purified using RNA Clean & Concentrator™-5 (Zymo Research Corp., R1015) before preparing small RNA libraries. Uniquely barcoded libraries of 2 ng PNK-treated RNA were prepared using NEXTFLEX® Small RNA-Seq Kit v3 (Perkin Elmer Inc., NOVA-5132-05) with 16 PCR cycles and no size selection protocol. The final libraries were quantified using the Agilent High Sensitivity DNA kit (Agilent Technologies, 5067-4626). The libraries were combined at 4 nM, denatured, and diluted to 1.5 pM using standard normalization protocol, and combined with diluted and denatured PhiX control before loading. Sequencing was performed using the NextSeq^®^ 500/550 Mid Output Kit v2 (300 cycles) with single indexing and paired-end reads.

#### Data processing and bioinformatics analysis

The raw RNA-seq data was trimmed using Cutadapt as per the manufacturer’s instructions (NEXTFLEX® Small RNA-Seq Kit v3) followed by quality analysis using FastQC software (*56*). Raw data (fastq files) were aligned to the human genome (GRCh38.p11) using Bowtie2-based sRNAbench to profile and map various RNA types and their respective counts (*57*). The countMA variable was used to create the count matrices for mRNAs and ncRNAs. The count matrices were then normalized and significantly differentially expressed mRNAs and ncRNAs were identified using *DESeq2* package in R (*58*). The p-value cutoff for statistically significant genes was 0.05 and log_2_FC cutoff for upregulated genes and downregulated genes was log_2_FC > 1 and log_2_FC < -1, respectively. Principal component analysis was conducted and plotted using *plotPCA* and *ggplot2* packages, respectively. Volcano plots were plotted using *EnhancedVolcano* package (v1.18.0) in R. Venn diagrams were adapted from the *ggvenn* package (v0.1.10) in R. Heatmaps were generated using *pheatmap* package (v1.0.12) in R with distance measure set to “manhattan” and clustering method set to “ward.D2”. Gene set enrichment and pathway analysis was performed using PANTHER (*59*, *60*). All R analysis was performed using statistically significant genes (*p* < 0.05) or gene ontology terms (p-adj < 0.05) except for principal component analysis and was conducted in RStudio environment (RStudio v2023.06.0 421 for Windows.

#### Correlation of EV-mRNA and EV-protein signature with Human Cell Atlas

The first-draft human cell atlas published by the Tabula Sapiens Consortium, which contains 1.1 million cells from 28 organs of 24 normal human subjects was used for our analysis (*61*). The lung cell atlas and global immune cell atlas were filtered to only include male individual. No such filtration was available for kidney cell atlas. In addition, to determine the disease specificity of EV-mRNA signature, the integrated Human Lung Cell Atlas Full was used, which consists of over 2 million cells from the respiratory tract of 486 individuals and includes 49 different datasets of which 35 datasets include donors with various lung diseases (*62*). Based on the pre-RESP vs. post-RESP comparison, two genesets were created that represented the top 50 EV-mRNAs in pre-RESP (pre-RESP^high^) or post-RESP (post-RESP^high^) samples. The mean expression of pre-RESP^high^ and post-RESP^high^ EV-mRNAs or mRNAs corresponding to pre-RESP^high^ and post-RESP^high^ EV-proteins across different cell types, tissue types, and disease types was visualized and compared using the CELLxGENE Explorer.

### EV cargo analysis of proteins using proteomics approaches

#### Sample preparation

The total protein concentration of the resuspended EV pellet from ultracentrifugation was determined using the Bicinchoninic acid (BCA) assay (Life Technologies, 23225). The volume of urine samples for mass spectrometry analysis was normalized based on protein concentration in EVs so that each sample contained an equal amount of injected total protein, and the EVs from urine were isolated using qEVoriginal (70 nm) size exclusion column (Izon Science Ltd.) and processed by Tymora Analytical Operations (West Lafayette, IN) using magnetic EVtrap beads as previously described (43). The isolated and dried EV samples were lysed to extract proteins using the phase-transfer surfactant (PTS) aided procedure (*63*). The proteins were reduced and alkylated by incubation in 10 mM tris(2-carboxyethyl)phosphine (TCEP) and 40 mM 2-chloroacetamide (CAA) for 10 min at 95°C. The samples were diluted five-fold with 50 mM triethylammonium bicarbonate and digested with Lys-C (Wako) at 1:100 (wt/wt) enzyme-to-protein ratio for 3 h at 37°C. Trypsin was added to a final 1:50 (wt/wt) enzyme-to-protein ratio for overnight digestion at 37°C. To remove the PTS surfactants from the samples, the samples were acidified with trifluoroacetic acid (TFA) to a final concentration of 1% TFA, and ethyl acetate solution was added at 1:1 ratio. The mixture was vortexed for 2 min and then centrifuged at 16,000 × g for 2 min to obtain aqueous and organic phases. The organic phase (top layer) was removed, and the aqueous phase was collected. This step was repeated once more. The samples were dried in a vacuum centrifuge and desalted using Top-Tip C18 tips (GlyGen) according to manufacturer’s instructions. The samples were dried completely in a vacuum centrifuge and stored at -80°C.

#### Data capturing and analysis

Dried peptide samples were dissolved in 4.8 μL of 0.25% formic acid with 3% (vol/vol) acetonitrile and 4 μL of each injected into an EASY-nLC 1000 (Thermo Fisher Scientific, Inc.). Peptides were separated on a 45-cm in-house packed column (360 μm OD×75 μm ID) containing C18 resin (2.2 μm, 100 Å; Michrom Bioresources, Inc.). The mobile phase buffer consisted of 0.1% formic acid in ultrapure water (buffer A) with an eluting buffer of 0.1% formic acid in 80% (vol/vol) acetonitrile (buffer B) run with a linear 60-min gradient of 6–30% buffer B at flow rate of 250 nL/min. The EASY-nLC 1000 was coupled online with a hybrid high-resolution LTQ-Orbitrap Velos Pro mass spectrometer (Thermo Fisher Scientific, Inc.). The mass spectrometer was operated in the data-dependent mode, in which a full-scan MS (from m/z 300 to 1,500 with the resolution of 30,000 at m/z 400), followed by MS/MS of the 10 most intense ions [normalized collision energy - 30%; automatic gain control (AGC) - 3E4, maximum injection time - 100 ms; 90sec exclusion].

#### Data processing

The raw files were searched directly against the human Swiss-Prot database with no redundant entries, using Sequest and Byonic search engine (Protein Metrics) loaded into Proteome Discoverer 2.2 software (Thermo Fisher Scientific, Inc.). Initial precursor mass tolerance was set at 10 ppm, the final tolerance was set at 6 ppm, and ion trap mass spectrometry (ITMS) MS/MS tolerance was set at 0.6 Da. Search criteria included a static carbamidomethylation of cysteines (+57.0214 Da), and variable modifications of oxidation (+15.9949 Da) on methionine residues and acetylation (+42.011 Da) at N terminus of proteins. Search was performed with full trypsin/P digestion and allowed a maximum of two missed cleavages on the peptides analyzed from the sequence database. The false-discovery rates of proteins and peptides were set at 0.01. All protein and peptide identifications were grouped, and any redundant entries were removed. Only unique peptides and unique master proteins were reported.

#### Label-free quantitation analysis

All data were quantified using the label-free quantitation node of Precursor Ions Quantifier through the Proteome Discoverer v2.2 (Thermo Fisher Scientific, Inc.). For the quantification of proteomic data, the intensities of peptides were extracted with initial precursor mass tolerance set at 10 ppm, minimum number of isotope peaks as 2, maximum ΔRT of isotope pattern multiplets – 0.2 min, PSM confidence FDR of 0.01, with hypothesis test of ANOVA, maximum RT shift of 5 min, pairwise ratio-based ratio calculation, and 100 as the maximum allowed fold change. For calculations of fold-change between the groups of proteins, total protein abundance values were added together, and the ratios of these sums were used to compare proteins within different samples.

### Statistical analysis

Statistically significant differentially expressed RNAs and proteins were determined using the respective RNAseq and proteomics data analysis pipeline. The statistical significance of ontology analyses was determined based on the default algorithm of the analysis tool. To compare the averages of two groups, unpaired t-test with Welch’s correction was used. Where applicable, results are reported as mean ± standard deviation. p < 0.05 was regarded as statistically significant.

## SUPPLEMENTARY MATERIALS

**Figure S1.** Summary of sequencing data of urinary EVs-RNA type, stratified by exposure status and time of urine sample collection. Pre, pre-respirator intervention (RI) samples, n=5. Post, post-RESP samples, n=5.

**Figure S2.** Mean expression level of EV-mRNA signature in lung, kidney and immune cell atlases.

**Figure S3.** Tissue-specific mean expression of pre-RESP^high^ EV-mRNA signature among specific cell types of myeloid leukocytes.

**Figure S4.** Tissue-specific mean expression of pre-RESP^high^ EV-mRNA signature among hematopoietic cells and neutrophils.

**Figure S5.** Mean expression level of mRNAs corresponding to pre-RESP-enriched (pre-RESP^high^) and post-RESP-enriched (post-RESP^high^) EV-proteins across different classes of cells in lung cell atlas.

**Table S1**. Summary of urine creatinine for pre-RESP and post-RESP worker samples.

**Data file S1**. Bioinformatics supplementary data.

## Supporting information

Bioinformatics supplementary data

## ACKNOWLEDGMENTS

The authors would like to especially thank the workers and management of the company for their collaboration with us and support of this work. The authors would also like to thank the Next Generation Sequencing, Genomics & Bioinformatics Lab at UMass-Lowell for their support in EV RNA isolation, Qubit and Agilent Bioanalyzer quantification of the isolated RNA, library preparation, and sequencing. The authors also extend their appreciation to Dr. Earl Ada and the Materials & Chemical Characterization Lab at UMass-Lowell for training and support with transmission electron microscopy. Additionally, the authors would like to thank Carolyn Marks and the Harvard Center for Nanoscale Systems for their support on cryo-electron microscopy.

## AUTHOR CONTRIBUTIONS

Conceptualization: IS, DB, AP

Methodology: IS, DB, JL

Software: IS

Formal analysis: IS, DB

Investigation: IS

Data Curation: IS

Visualization: IS

Validation: IS, PL

Funding acquisition: IS, DB

Project administration: IS, DB

Supervision: IS, DB

Writing – original draft: IS, DB

Writing – review & editing: IS, DB, AW, CR

## DATA AND MATERIALS AVAILABILITY

All data generated or analyzed during this study are included in this published article (and its Supplementary Information files). The raw files generated during the current study are available from the corresponding author on reasonable request.

