## Supplementary Materials for "A cross-week analysis of urinary extracellular vesicles after respiratory tract exposure intervention identifies systemic signaling changes – a pilot study of isocyanate-exposed workers"

SUPPLEMNTARY MATERIALS

^3^ Next Generation Sequencing & Genomics Lab

UMass Lowell Core Research Facilities

Lowell, MA 01854

^4^ Department of Chemistry

University of Massachusetts Lowell

Lowell, MA 01854, USA

^5^Yale University

Department of Internal Medicine,

Yale University School of Medicine

New Haven, CT 06510, USA

^6^Department of Biomedical and Nutritional Sciences, Zuckerberg College of Health Sciences

University of Massachusetts Lowell

Lowell, MA 01854, USA

*Corresponding authors.

Ikjot S. Sohal –

Dhimiter Bello –

**RESULTS**

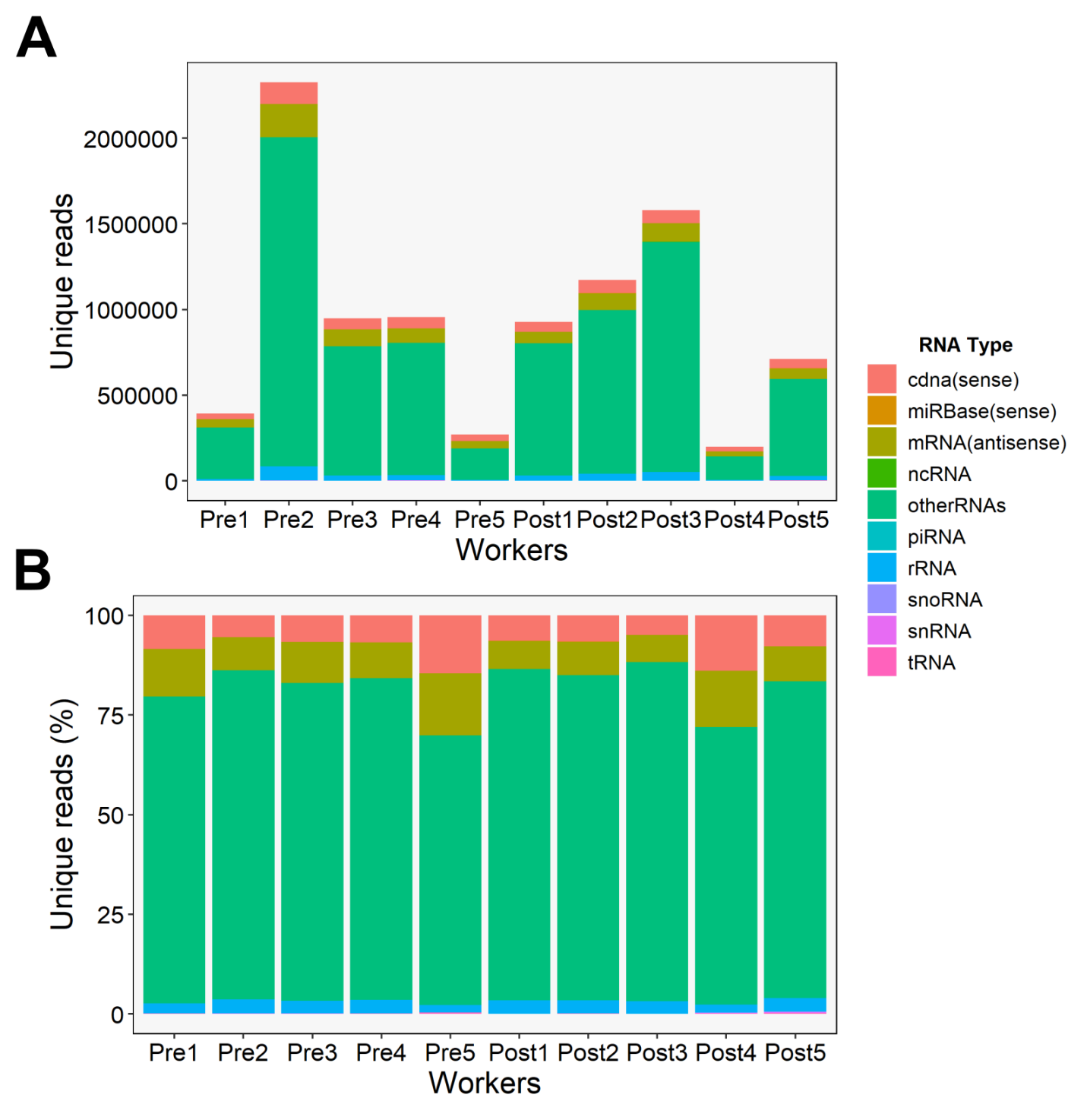

**Figure S1**. Summary of sequencing data of urinary EVs-RNA type, stratified by exposure status and time of urine sample collection. Pre, pre-respirator intervention samples, n=5. Post, post-respirator intervention samples, n=5.

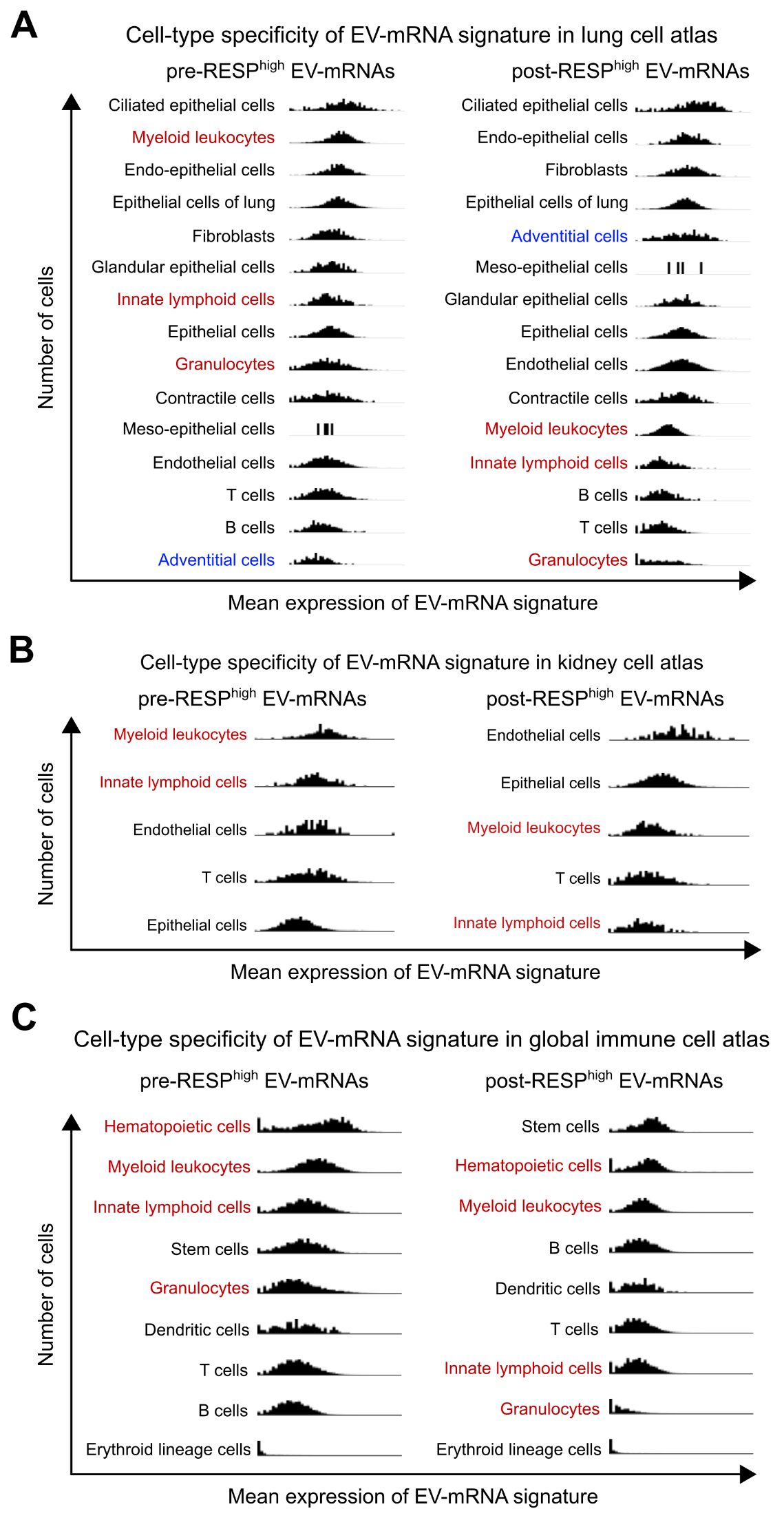
**Figure S2. Mean expression level of EV-mRNA signature in lung, kidney and immune cell atlases**. (**A**) Mean expression of pre-RESP-enriched (pre-RESP^high^) and post-RESP-enriched (post-RESP^high^) EV-mRNAs across different classes of cells in lung cell atlas, sorted by highest to lowest mean expression. Cell types highlighted in red indicate higher expression of pre-RESP^high^ EV-mRNAs and in blue indicate higher expression of post-RESP^high^ EV-mRNAs. (**B**) Mean expression of pre-RESP^high^ and post-RESP^high^ EV-mRNAs across different classes of cells in kidney cell atlas, sorted by highest to lowest mean expression. Cell types highlighted in red indicate higher expression of pre-RESP^high^ EV-mRNAs. (**C**) Mean expression of pre-RESP^high^ and post-RESP^high^ EV-mRNAs across different classes of cells in global immune cell atlas, sorted by highest to lowest mean expression. Cell types highlighted in red indicate higher expression of pre-RESP^high^ EV-mRNAs and in blue indicate higher expression of post-RESP^high^ EV-mRNAs.

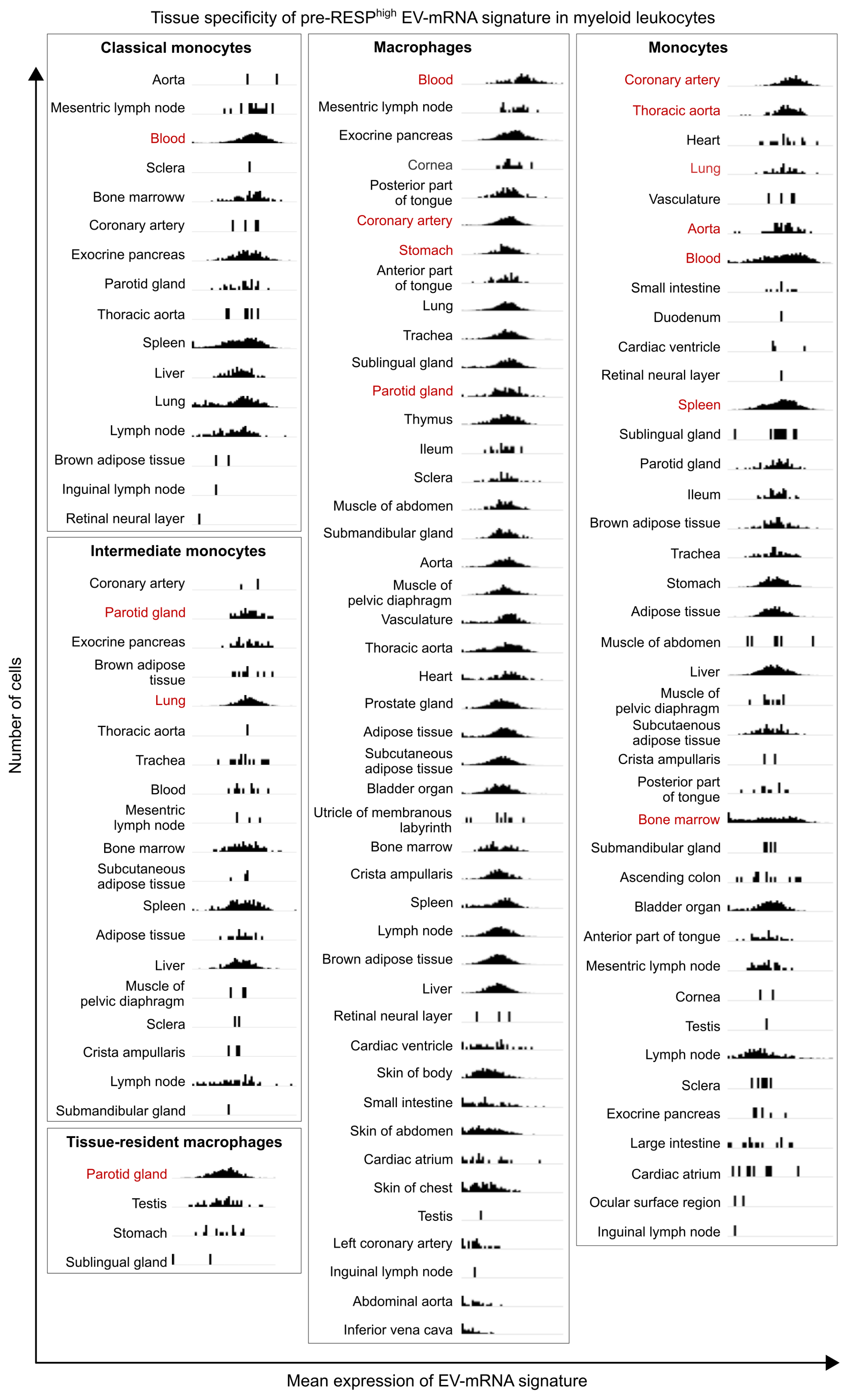
**Figure S3.** Tissue-specific mean expression of pre-RESP^high^ EV-mRNA signature among specific cell types of myeloid leukocytes – classical monocytes, macrophages, monocytes, intermediate monocytes, and tissue-resident macrophages, sorted by highest to lowest mean expression. Tissue types highlighted in red indicate higher expression of pre-RESP^high^ EV-mRNAs.

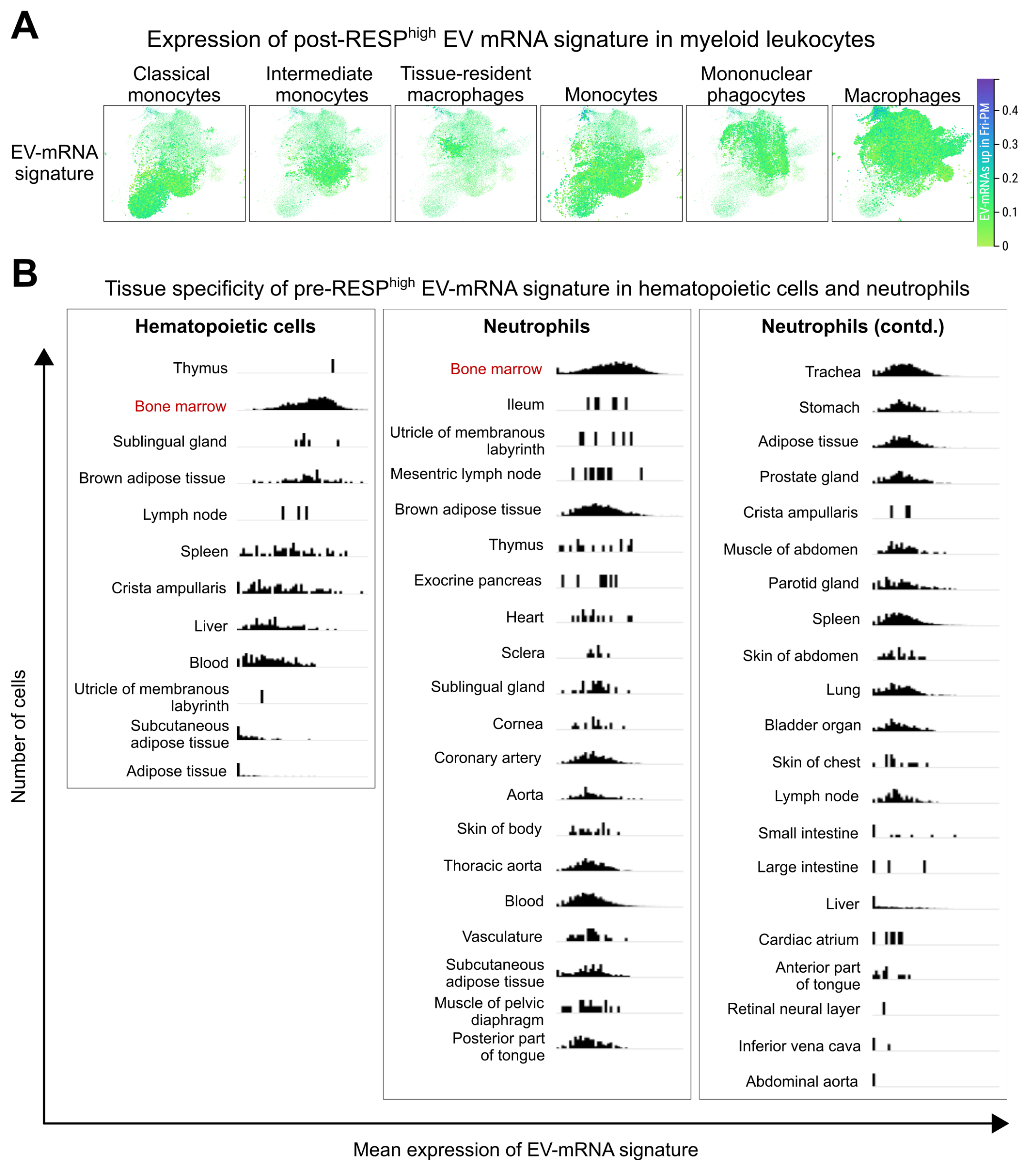
**Figure S4. Expression in myeloid leukocytes, hematopoietic cells and neutrophils**. (**A**) UMAP-based plot demonstrating the expression of post-RESP^high^ EV-mRNAs in specific cell types among myeloid leukocytes. Color bar indicates expression level. (**B**) Tissue-specific mean expression of pre-RESP^high^ EV-mRNA signature among hematopoietic cells and neutrophils, sorted by highest to lowest mean expression. Tissue types highlighted in red indicate higher expression of pre-RESP^high^ EV-mRNAs.

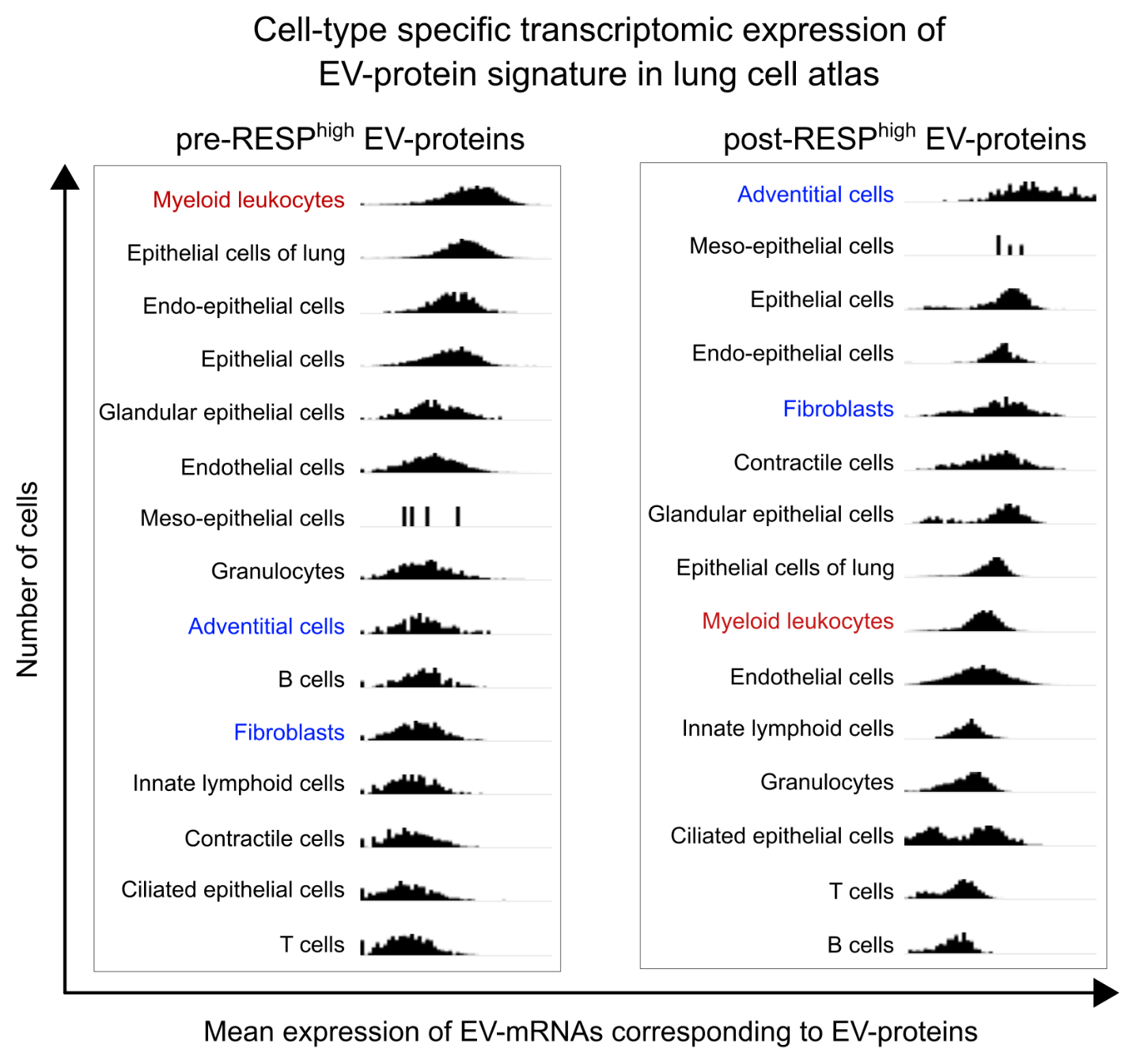
**Figure S5.** Mean expression level of mRNAs corresponding to pre-RESP-enriched (pre-RESP^high^) and post-RESP-enriched (post-RESP^high^) EV-proteins across different classes of cells in lung cell atlas, sorted by highest to lowest mean expression. Cell types highlighted in red indicate higher expression of mRNAs corresponding to pre-RESP^high^ EV-mRNAs and in blue indicate higher expression of mRNAs corresponding to post-RESP^high^ EV-mRNAs.

**Table S1**. Summary of urine creatinine for pre-RESP and post-RESP worker samples.

| **Category** | **Subject #** | **Unnormalized EV concentration** | **Creatinine concentration (mmol/L)** | **Normalized EV concentration** | **Average normalized EV concentration** |
| --- | --- | --- | --- | --- | --- |
|  |  | #/mL (× 10^10^) | mmol/L | #/mmol creatinine (× 10^12^) | Mean (Min, Max) |
| pre-RESP | M1 | 11.90 | 20.69 | 5.75 | 3.42 (0.91, 8.41) |
|  | M2 | 6.83 | 30.49 | 2.24 |  |
|  | M3 | 7.06 | 14.96 | 4.72 |  |
|  | M4 | 2.53 | 27.70 | 0.91 |  |
|  | M5 | 4.93 | 5.86 | 8.41 |  |
| post-RESP | F1 | 4.92 | 53.90 | 0.91 | 5.56 (0.91, 25.56) |
|  | F2 | 9.80 | 25.40 | 3.86 |  |
|  | F3 | 20.40 | 19.59 | 10.42 |  |
|  | F4 | 11.30 | 20.02 | 5.64 |  |
|  | F5 | 21.00 | 8.22 | 25.56 |  |
